# Multicentric tracking of multiple agents by anterior cingulate cortex during pursuit and evasion

**DOI:** 10.1101/796375

**Authors:** Seng Bum Michael Yoo, Jiaxin Cindy Tu, Benjamin Yost Hayden

## Abstract

Successful pursuit and evasion require rapid and precise coordination of navigation with adaptive motor control. We hypothesized that the dorsal anterior cingulate cortex (dACC), which communicates bidirectionally with both the hippocampal complex and premotor/motor areas, would serve a mapping role in this process. We recorded responses of dACC ensembles in two macaques performing a joystick-controlled continuous pursuit/evasion task. We found that dACC multiplexes two sets of signals, (1) world-centric variables that together form a representation of the position and velocity of all relevant agents (self, prey, and predator) in the virtual world, and (2) avatar-centric variables, i.e. self-prey distance and angle. Both sets of variables are multiplexed within an overlapping set of neurons. Our results suggest that dACC may contribute to pursuit and evasion by computing and continuously updating a multicentric representation of the unfolding task state, and support the hypothesis that it plays a high-level abstract role in the control of behavior.

## INTRODUCTION

Foragers often encounter mobile prey that are capable of fleeing them. Not surprisingly, pursuit is a major element of the behavioral repertoires of many foragers [1–5]. Likewise, many foragers must also avoid predators seeking to capture them (e.g. 6). Recent studies have begun to identify the computational processes underlying pursuit and evasion behavior (hereafter shortened to pursuit) in several species [7,8]; nonetheless, the neural bases of these behaviors remain almost wholly unexplored (but see 9). Despite this relative paucity of scholarly interest, pursuit is an important problem in neuroscience because it is common in mobile animals, and because it is highly determinative of reproductive success and thus a likely driver of evolution. Moreover, it represents a mathematically tractable form of continuous decision-making, which decision neuroscience often ignores in favor of discrete decisions [10,11].

When foragers move in their environments, neurons in the hippocampus and adjacent structures track the forager’s own positions of using a firing rate code [12–14]. They do so by means of an explicit cognitive map [15,16]. Specifically, each neuron in the hippocampus or medial entorhinal cortex exhibits one or more preferred firing fields [13]. That is to say, entry by an animal into a specific location results in robust spiking activity. The hippocampal spatial map is allocentric, meaning that it is organized relative to external space [12,17]. To employ this information to guide actions, however, foragers must use a complementary egocentric coding system, that is, one that is relative to the self [17–19]. Egocentric spatial representations are related to action planning and are often associated with the premotor cortex/primary motor cortex [20,21] and sometimes with the parietal and posteromedial cortex [21,22]. Even when navigating virtual or abstract environments, foragers can benefit from multiple reference frames. That is, they can make an abstract, or *world-centric* representation, but when the time comes to perform an action, or to navigate the virtual space, they may need to use a coordinate system that is aligned to the framework of their response modality.

In addition to monitoring one’s place in space, pursuit requires the careful coordination of two distinct processes: (1) the computation and dynamic updating of a representation of the pursuit environment, including the kinematics of the prey and predator; (2) the ability to select and quickly adjust behavior in response to changing demands. In other words, pursuit requires the coordination of cognitive mapping functions with motor control functions. To understand the cognitive mapping element of this process, we were especially interested in brain regions that have strong inputs from, on one hand, the hippocampal complex and, on the other, the premotor and motor system. The dorsal anterior cingulate cortex (dACC) fits this description [23–26]. Specifically, it is one of a small number of regions that receive converging information from reward regions (in this case, orbitofrontal cortex, amygdala, and insula) and navigational brain regions (parahippocampal and entorhinal cortices, and the hippocampal formation), and provides a direct output to motor brain regions, including the primary and supplementary motor cortices.

Some evidence supports the idea that both rodent and human dACC may carry place-relevant information, suggesting it may play a mapping role in pursuit [27–31]. We hypothesized dACC carry a rich and dynamic representation of key variables needed for pursuit decisions. Note that there is no *a priori* reason to assume that pursuit and evasion both rely on the same circuits. Indeed, it is likely that the neuroanatomy that mediates these two processes differs at least somewhat. Still, we reasoned that they may have some overlap and that this overlap, if it exists, is may be associated with the dACC.

We recorded responses of neurons in the dACC of two macaques performing a real-time pursuit task. Subjects used a joystick to smoothly and rapidly move an avatar around a virtual pen on a computer screen to pursue fleeing prey and avoid predators that were chasing them. All agents other than the subject were controlled by interactive algorithms that used video game-derived artificial intelligence (AI) strategies. We found that dACC neurons track world-centric kinematic variables (specifically, position, velocity, and acceleration) for all three agents (self, prey, and predator). Although the responses of dACC neurons are spatially selective, they are more complex and multimodal than a place or grid cells would be (and in this, they resemble non-grid cells of the medial entorhinal cortex, ref 32). Neurons in dACC also track the two key avatar-centric variables - relative position and angle of the self. Together, these results highlight the sophisticated role of dACC in monitoring complex relational positions and provide a basis for understanding the neuroscience of pursuit and evasion.

## RESULTS

### Pursuit and evasion behavior of macaques

We measured responses from macaque dACC neuronal ensembles collected during a demanding computerized real-time *pursuit task* (subject K: 5594 trials; subject H: 2845 trials, **Methods**). A subset of these data was analyzed and summarized for a different study; all results presented here are new [33]. On each trial, subjects used a joystick to control the position of an avatar (a yellow or purple circle) moving smoothly in a rectangular field on a computer monitor (Figure 1A-D and **Supplementary Video**). Capture of prey (a fleeing colored square) yielded a juice reward delivered to the subject’s mouth via a metal tube. The prey item on every trial was drawn randomly from a set of five that differed in maximum velocity and associated reward size. On 50% of trials (randomly determined) subjects had the opportunity to pursue either or both of two different prey items (but could only capture one). On 25% of trials (randomly determined), subjects also had to avoid one of five predators (a pursuing colored triangle). Capture by the predator ended the trial early, imposed a timeout penalty, and resulted in no reward.

**Figure 1.**
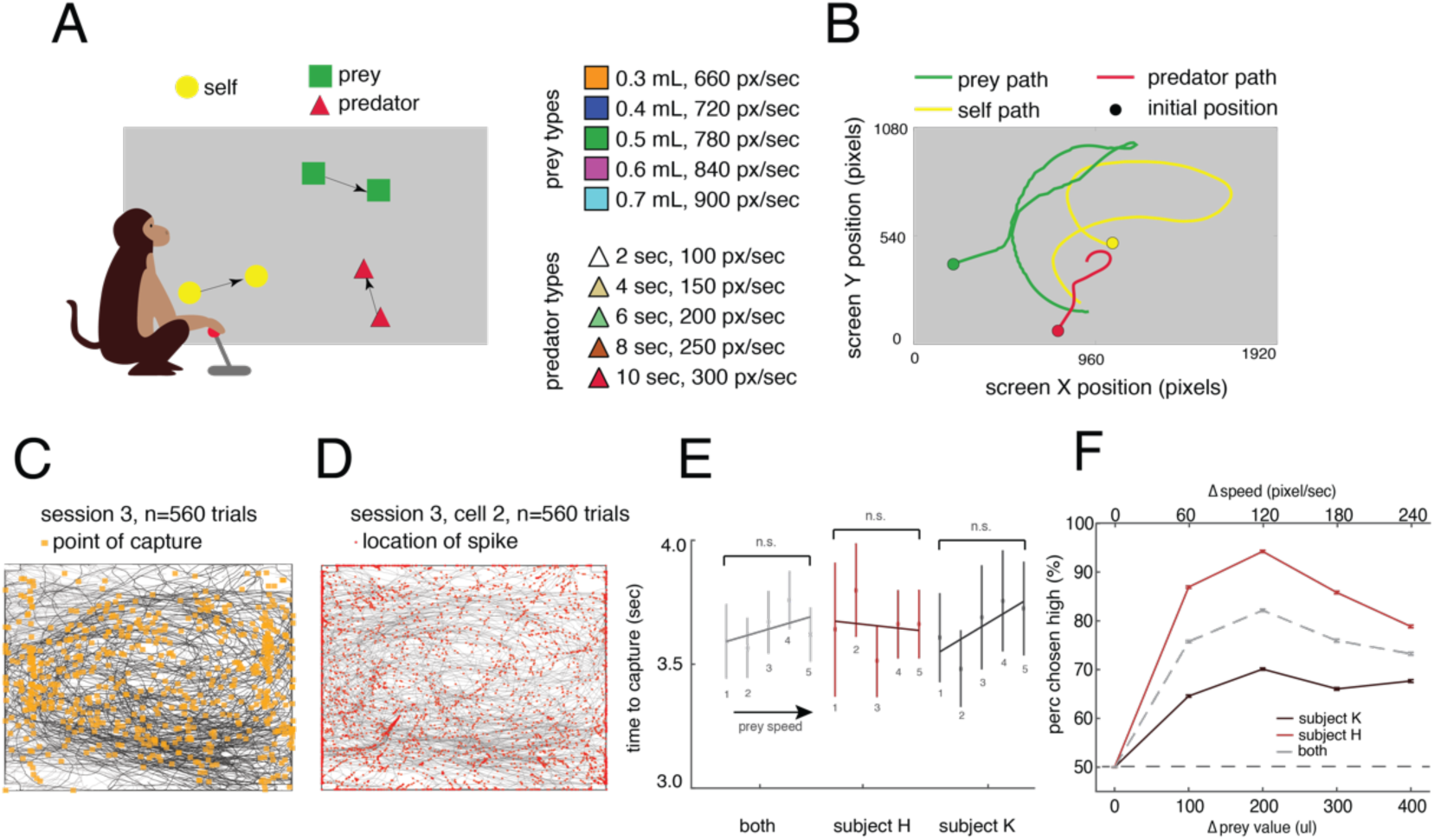
Experimental Paradigm and Behavior Results. **(A)** Cartoon of the *virtual pursuit task*. Subject uses a joystick to control an avatar (circle) to pursue prey (square) and avoid predators (triangle). **(B)** Agent trajectories on example trial. **(C)** Illustration of typical session indicating subject’s path (black lines) and points of capture (orange dots). **(D)** Illustration of example session showing subjects’ path (gray lines) and points of spiking (red dots). **(E)** Average time to capture for each subject as a function of value of prey. No significant relationships are observed despite large datasets. This lack of correlation suggests that subjects generally traded off effort to maintain roughly constant pursuit duration. **(F)** The proportion of trials on which subjects chose the larger value prey item despite it being faster was greater than 50% for each prey difference (other than matched). Error bars indicate the standard error of the mean.

Subjects successfully captured the prey in around 80% of trials (subject K: 78.95%; subject H:84.91%). The average time for capturing a prey was 3.85 seconds (subject K: 4.05; subject H: 3.50; Figure 1E). Capture time did not differ according to the prey value/speed (F=50.98, p=0.3797 for subject K, F=26.68, p=0.6118 for subject H, two-way ANOVA). On trials in which subjects faced two prey (50% of all trials), they had to choose which to pursue. On these trials, subjects chose the higher valued prey more often, even though they were faster and presumably more difficult to catch (K: 67.10%, H: 86.46%, Figure 1F). Notably, whenever the subject was faced with two prey that differed in size, they reliably chose the one with the larger value (and thus the one with the faster speed as well). These patterns suggest that subjects understood that prey color provided valid information about the value and/or speed of the prey and used this information to guide behavior. This pattern also suggests that, for the parameters we chose, the marginal increase in reward value was more effective at influencing choice, on average, than the marginal increase in capture difficulty.

### World-centric encoding in dACC

We recorded neural activity during performance of the task (n=167 neurons; 119 in subject H and 48 in subject K). We applied a Generalized Linear Model approach (GLM, refs 32,34–36) based on a Linear-Nonlinear (LN) model that does not assume any parametric shape of the tuning surface (see ref 32, Figure 2A, B). This procedure includes a cross-validation step, meaning that the results are essentially validated for statistical significance against a randomized version of the same data. This approach effectively includes a test for reliability, and also efficiently uses information about spatial coherence, to detect significant spatial selectivity. Note that, although we don’t report the data, we confirmed that all results presented below are observed in both subjects individually, except in one case (predator angle in subject K).

**Figure. 2.**
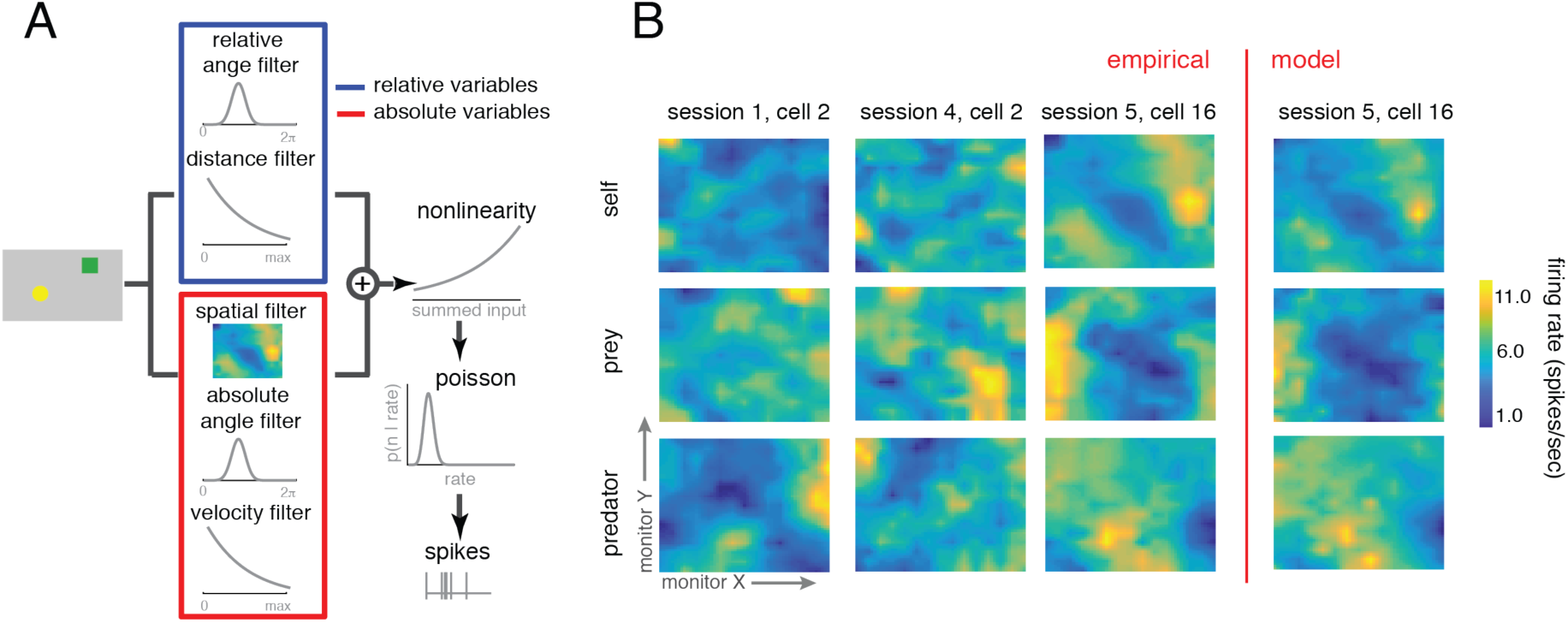
Examples of multi-agent world-centric mapping functions in dACC. **(A)** Schematic of the analysis approach we took, see **Methods** and ref 32. **(B)** Maps of the responses of three example neurons with significant spatial selectivity, chosen to illustrate the typical types of responses we see. X- and Y-dimensions in each subplot correspond to X- and Y-dimensions of the computer monitor; the Z-dimension corresponds to the firing rate of that cell. Yellower areas indicate spots in which entry tended to result in spiking. Top row: responses to self-position. Middle row: responses to prey position. Bottom row: responses to predator position. Rightmost column: responses of the model (see **Methods**) for the same neuron whose observed responses are shown in the adjacent column.

Our analysis approach is a way of asking whether a neuron shows significant tuning (for example, whether it has angular tuning, Figure 3A) but is agnostic about the shape of the tuning (for example, whether that place field is localized to a point, as hippocampal place fields are, or has a more complex shape). To identify the simplest model that best described neural spiking, we used a forward-search procedure that determined whether adding variables significantly improved model performance with 10-fold cross-validation (Figure 3 B-D).

**Figure 3.**
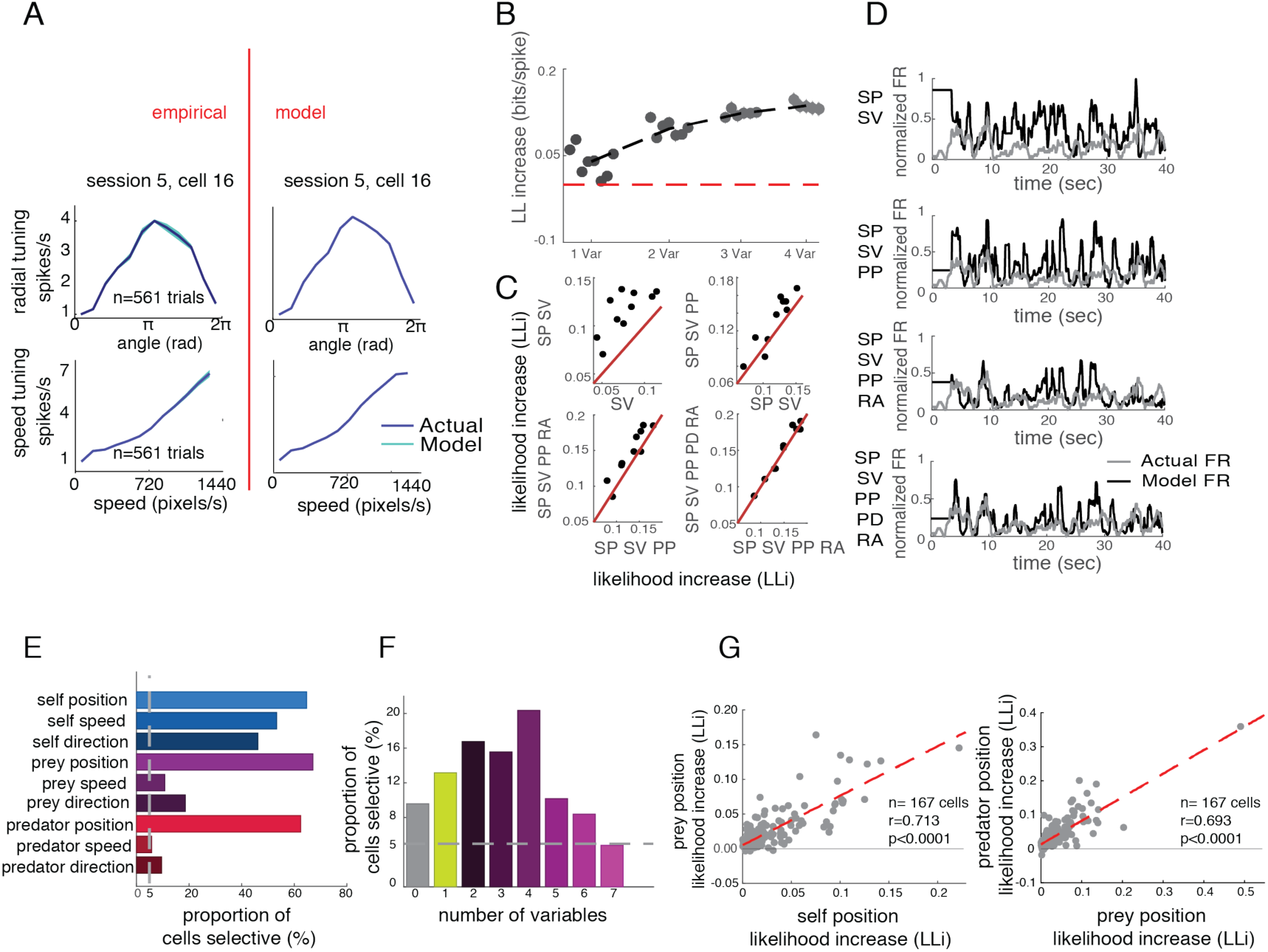
World-centric tracking in dACC neurons. **(A)** Direction and speed tuning (left) and model fit (right) for an example neuron. **(B)** Proportion of neurons selective for 1, 2, 3, or 4 of the world-centric variables self position, self velocity, prey position, and relative angle between self and prey. Horizontal jitter added for clarity. Red dashed horizontal line indicates the likelihood value for mean firing rate model, which we treated as a zero variable model. **(C)** Likelihood increase in fit as a function of adding additional variables. This method was used to select variables via cross-validation. The significance of tuning with respect to the variable was determined by sign-rank test between red unity line and 10-fold cross-validation data points. **(D)** Example neuron showing observed firing rate (gray) and model predicted firing rate (black). **(E)** Proportion of neurons tuned for each possible number of task variables. Proportions do not sum to 1 because neurons may be tuned for multiple variables; results reflect correction for multiple comparisons. **(F)** Proportion of neurons in our sample selective for any of 7 number of variables. Most neurons are selective for multiple of the variables we test. **(G)** Correlation between log-likelihood increase (LLi) for self-position vs. prey-position (left) and prey-position vs. predator-position (right). Each dot corresponds to one neuron. Positive correlation indicates selectivity is a shared property of selective cells.

For neural analyses, we focused on the *whole trial epoch*, that is, the period from the time when all agents appear on the screen (trial start) until the end of the trial, defined as either (i) the time when the subject captures the prey, (ii) the time when the predator captures the subject, or (iii) 20 seconds pass without either other event occurring. Examining this epoch, we found that 90.4% (n=151/167) of neurons are task-driven, meaning that neuronal responses depend on one or more of the variables we tested (Figure 3A, B, E, F). Note that the structure of our analysis, which forward searches for tuning for each variable automatically corrects for multiple comparisons (i.e. equivalent to using a cutoff of p=0.05, corrected). The majority of neurons showed sensitivity to the spatial position of the avatar (64.5%, n=108/167). Roughly similar proportions of cells showed sensitivity to the position of the prey (65.5%, n=112/167), and to the position of the predator (59.3%, n=89/150). We found that responses of 22.0% (n = 33/150) of neurons are selective for the positions of all three agents. Note the predator fits were done separately because they only occurred on 25% of trials, and for that analysis, we removed 17 cells that had trial counts below our *a priori* threshold. Note also that there is no certainty that subjects are engaging in pursuit and evasion simultaneously; indeed, it may well be the case that they alternate between these two modes.

We next tested whether overlapping populations of neurons encode self and prey position by examining log-likelihood increase (LLi) associated with adding the relevant variables (Figure 3G). For each variable pair, we found a positive LLi relationship, indicating that neurons encoding one variable are more likely to encode the other, and therefore, evidence against specialized subpopulations of neurons for these variables. In other words, we found that populations overlap more than expected by chance (self/prey r=0.7882; self/predator: r=0.7092; prey/predator: r =0.6548; p<0.001 for all cases, Pearson correlation). This finding indicates that coding strength is positively correlated for each pair and that the coding comes from a highly overlapping set of populations rather than from distinct subpopulations (see ref 37 for motivation for this analysis approach). Thus, the two populations of neurons overlap more than might be expected by chance if these effects were distributed at random in the population. This result thus allows us to reject the hypothesis that the two groups of neurons come from distinct sets - or even from overlapping sets that diverge more than might be expected by chance. Nonetheless, while these results are consistent with the idea that the neurons come from a single population, they are also consistent with the idea that they come from populations that overlap more than chance but are still partially distinct.

We next used a previously published method to assess how the spatial kernels for the three agents compare (spatial efficiency or SPAEF, see ref 38). For each pair of agents, we focused on neurons that show significant tuning for both agents individually. These groups consisted of, respectively, subject *and* prey 24.0%, n = 36/150; subject *and* predator: 26.0%, n=39/150; prey *and* predator: 39.3%, n=59/150). Incidentally, the largest of these three variables, perhaps surprisingly, was for the prey-predator. It’s not clear why this is. One possibility is that this variable was encoded most strongly because of the special difficulty subject face in coordinating between pursuit and evasion strategies, and the need to attend to both other elements when doing so. Future work will be needed to test this hypothesis.

SPAEF is more robust than simple pairwise correlation because it combines three measures into a single value. Specifically, it combines pairwise correlation, coefficient of variation of spatial variability, and intersection between observed histogram and simulated histogram, see **Methods**). Across all neurons, we found that the SPAEF value between the subject and prey was −0.3282. This negative value indicates that the kernels are anti-correlated – locations that led to enhanced firing when the subject entered led to reduced firing at times when the prey entered it. This SPAEF value is significantly less than zero (p<0.001, Wilcoxon sign rank test). The value for the subject and predator was −0.2463. The analogous value for the prey and the predator was −0.2927. Both of these are also less than zero as well (p<0.001, Wilcoxon sign rank test). These findings indicate that neurons use distinct – and to some extent, anti-correlated - spatial codes for tracking the positions of the three agents. Consequently, these results suggest that dACC carries sufficient information for decoders to estimate path variables for all three agents.

We next asked whether a substantial number of neurons encode “self vs. other”. To do this, we examined the set of neurons with significant selectivity for self-position and for prey *and/or* predator position, and that showed a high positive predator-prey SPAEF value (that is, did not distinguish prey from predator). We found that 6 neurons meet these criteria (mean SPAEF value among those neurons is 0.2501). This proportion (3.6% of cells) is not significantly different from chance, suggesting that self vs. other encoding is not a major factor driving dACC responses, and that dACC differentiates prey from predator.

We next sought to characterize the size of these effects. To do so, we first selected neurons that showed significant selectivity for the position of each agent. Then, for each neuron in each set, we selected the peak firing rate and lowest firing rate in the 2D space. This measure is analogous to peak-to-trough measures for any other task variable. To assess population measures, we then computed the median within each set. (Median is more conservative than mean; it more effectively excludes outlier measurements). We find substantial effects for each category: self-tuned neurons: 13.34 spike/sec (95% confidence interval: 9.44 to 17.24 spike/sec); prey-tuned neurons: 11.77 spike/sec (95% confidence interval: 7.65 to 15.89 spike/sec); predator-tuned neurons: 12.55 spike/sec (95% confidence interval: 7.21 to 17.09 spike/sec). These effects are quite robust, and are comparable to modulations associated other factors in more conventional laboratory tasks.

### Speed information is also processed in dorsal anterior cingulate cortex

We hypothesized that dACC would encode agent speed. To test that idea, we added speed filters in our GLM and fit against the neural data. We found that 22.7% (n=34/150) neurons are selective for the speed of the self, 10.0% (n=15/150) of neurons for the speed of the prey, and 10.0% (n=15/150) for the speed of the predator (Figure 3A). Naturalistic tasks such as ours provide the opportunity to understand higher dimensional tuning than other methods. To gain insight into the diversity of speed tuning profiles, we performed an unsupervised k-means clustering on speed filters across the agents (Figure 4). Initially, we performed principal component analysis (PCA) on the filter coefficients. Then we obtained eigenvector of top 2-dimension in which explained more than 70% of variance of the data. We find both monotonically and ditonic speed filters (11 neurons for cluster 1, 16 neurons for cluster 3). Previous literature suggested that better perceptual discrimination for lower speed [39]. Interestingly, our result shows that neurons in ACC, at least, is not biased towards representing low speed but exhibit diversity.

**Figure 4.**
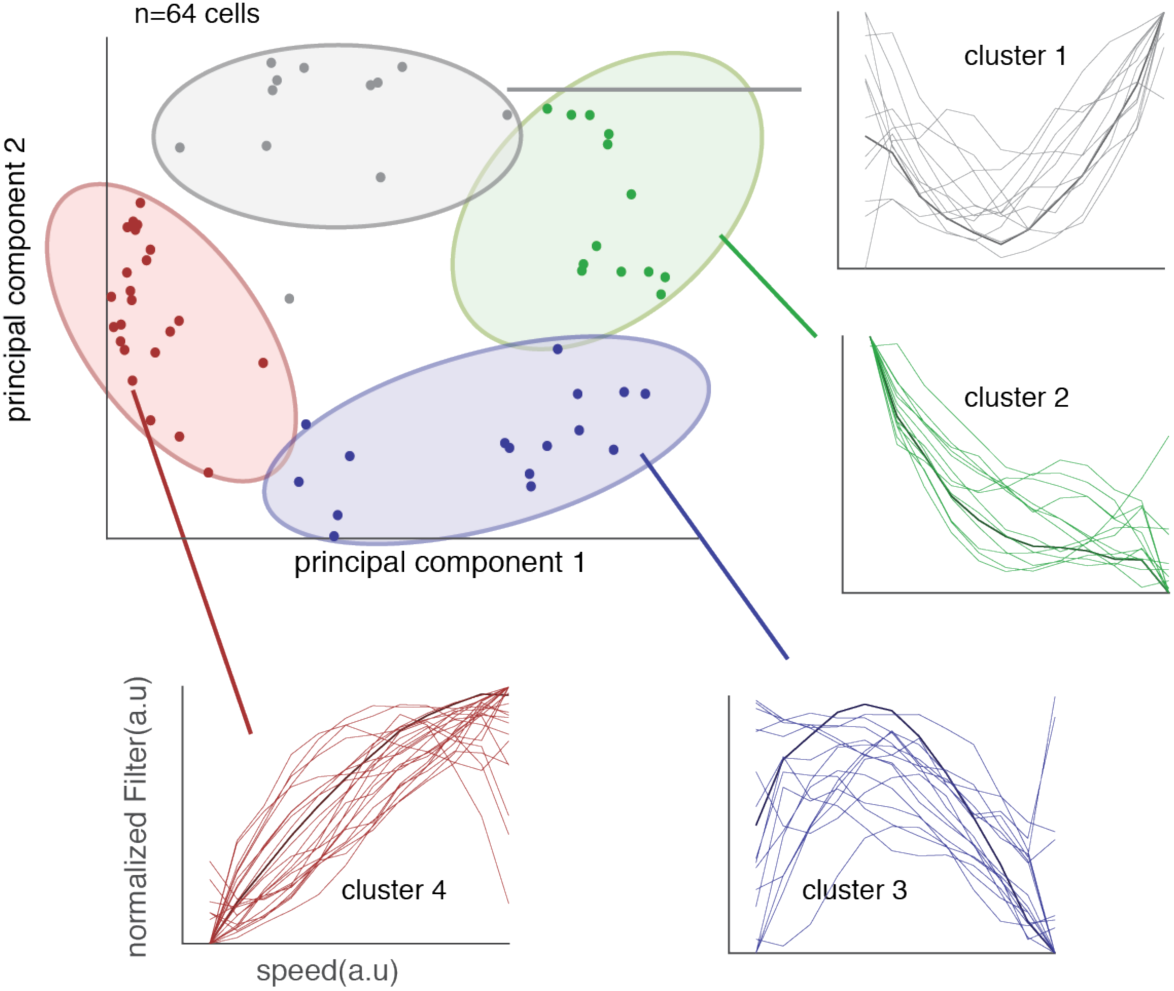
Diversity in responses of neurons to speed. We examined whether speed tuning reflects a single response profile, as would be expected if, for example, speed effects were simply an artifact of arousal. We performed a PCA procedure on all tuning curves and plot the results as a function of each PC. We then cluster the resulting patterns and show all tuning curves within each cluster. The diversity of responses, and especially the existence of clearly ditonic clusters (clusters 1 and 3), argues against an arousal confound.

### Avatar-centric encoding

We next examined avatar-centric coding, that is, coding of the position of the prey relative to the agent. According to our GLM, 37.7% of neurons (n=63/167) in our sample encode the distance between self and prey and 25.1% of neurons (n=42/167) encode the angle between self and prey (Figure 5A). Together, these two variables define the entire basis set of avatar-centric spatial variables relevant to the pursuit of the prey. That is, other avatar-centric variable can be expressed as a linear combination of them, and thus are available to decoders that have access to the responses of these neurons. A smaller proportion of neurons signal these variables relative to the predator (n=14/150, 9.3% for relative distance to predator, n=8/150, 5.3% for relative angle to predator). Note that the value for relative distance to predator is significant while the value for angle is not (distance: p=0.0220; angle: p=0.8496; one-way binomial test; Figure 5B, C). Nonetheless, *in toto*, these results indicate that dACC neurons carry a rich representation of the avatar-centric world in this task.

**Figure 5.**
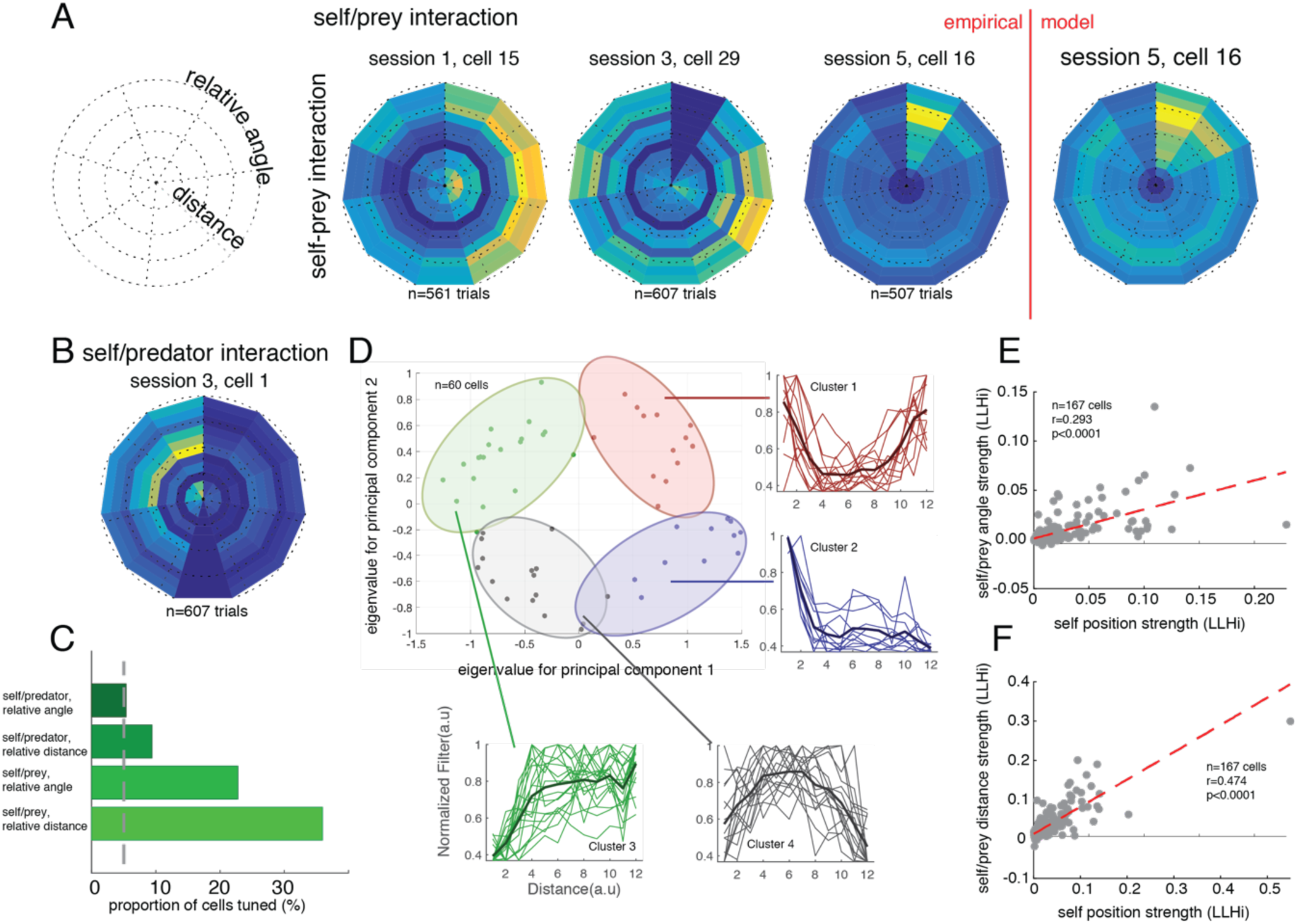
Avatar-centric Tuning in dACC. **(A)** Three example neurons and model fit for self-prey avatar-centric variables (angle and distance). Each radial plot shows the responses of a single neuron to prey located at several distances (radial dimension) and angles (angular dimension) relative to the agent’s avatar. Horizontal-right direction (0 degrees) reflects distance in front of the direction the avatar is currently travelling; horizontal left (180 degrees) indicates travel away from the prey. **(B)** Example neuron tuned for self-predator avatar-centric variables using same conventions as in panel A. **(C)** Proportion of neurons tuned for each of four avatar-centric variables. **(D)** Clusters of relative distance tuning functions (conventions as in Figure 4). Scattergram reflects results of principal components analysis (PCA). Clusters 3 and 4 are ditonic; their existence argues against low-level explanation in terms of arousal or reward proximity. **(E-F)** Correlation between log-likelihood increase (LLi) for self-position vs. self/prey distance (E) and angle (F). Each dot corresponds to one neuron. Positive correlation indicates that neurons selective for one variable tend to be more selective for another one. That in turn implies that tuning for the two variables comes from a single larger population of cells (or from highly overlapping populations) rather than distinct populations.

We were concerned that distance tuning, as determined by this analysis, may be artifactual - it may reflect proximity to reward, which is known to consistently enhance activity in dACC [40–44]. To test this alternative hypothesis, we performed an analysis of the diversity of responding. Specifically, we reasoned that if neurons encode distance, they will show a heterogeneity in response patterns but if they encode proximity to reward, they will show a more homogeneous and positive-going pattern. To examine our hypothesis, we clustered the shape of subject-prey distance filters (Figure 5D). These figures use the following radial plot conventions. The angle on the plot relative to 0 (i.e. horizontal and to the right) reflects the angle between the subject’s own avatar and the prey. Thus, a neuron selective for the subject bearing directly towards the avatar will have lighter colors on the right hand side of the radial plot. The radial dimension on the plot indicates the distance - thus, a neuron selective for distant prey will have lighter colors on the outer ring of the plot.

We observed a heterogeneity of curves, including a substantial fraction of neurons with decreasing and even ditonic curves (48.3%, n = 29/60, p < 0.001, two-way binomial tests). The ditonicity (i.e. positive and negative slopes within a single curve) of some neurons is important – it indicates that these neurons do not simply exhibit ramping behavior. This result thus argues against the possibility of avatar-centric distance simply being an artifact of the proximity of reward and/or arousal or other low-level features that scale with distance to reward (Figure 5D).

By identifying avatar-centric-coding neurons, we were able to ascertain whether avatar- and world-centric coding neurons arose from different or similar populations. We used the same log-likelihood correlation approach described above. We find that they are not distinct; instead, they overlapped considerably more than would be expected by chance (that is, the correlation of log-likelihoods was greater than zero, r=0.295, p<0.001; Figure 5E, F). This result is consistent with the possibility that these neurons come from a single task-selective population as well as with the possibility that they come from highly overlapping but partly distinct sets.

### Mixed selectivity

Encoded variables interacting non-linearly (mixed selectivity) is potentially diagnostic of control processes and can be harnessed for flexible responding [32,44–46]. We used two methods to test for mixed selectivity (**Methods** and Figure S3). First, we computed direction and speed tuning separately in high and low firing rate conditions (a method found in ref 46, Figure 6A). Then we performed regressions for the two conditions separately. A slope different from 1 indicates a multiplicative shift; an offset different from 0 indicates an additive shift. In our data, the median of the slope was significantly less than 1 (median slope=0.8533, p<0.022, rank-sum test) with little evidence for additive modulation (position: median bias=0.4909, p<0.001, rank-sum test; speed: median multiplicative factor=0.7692, p=0.048; median additive factor=0.4661, p<0.001; Figure 6B, C). This result indicates that dACC ensembles have the capacity to represent information in high-dimensional space by encoding multiple variables nonlinearly [45,48].

**Figure 6.**
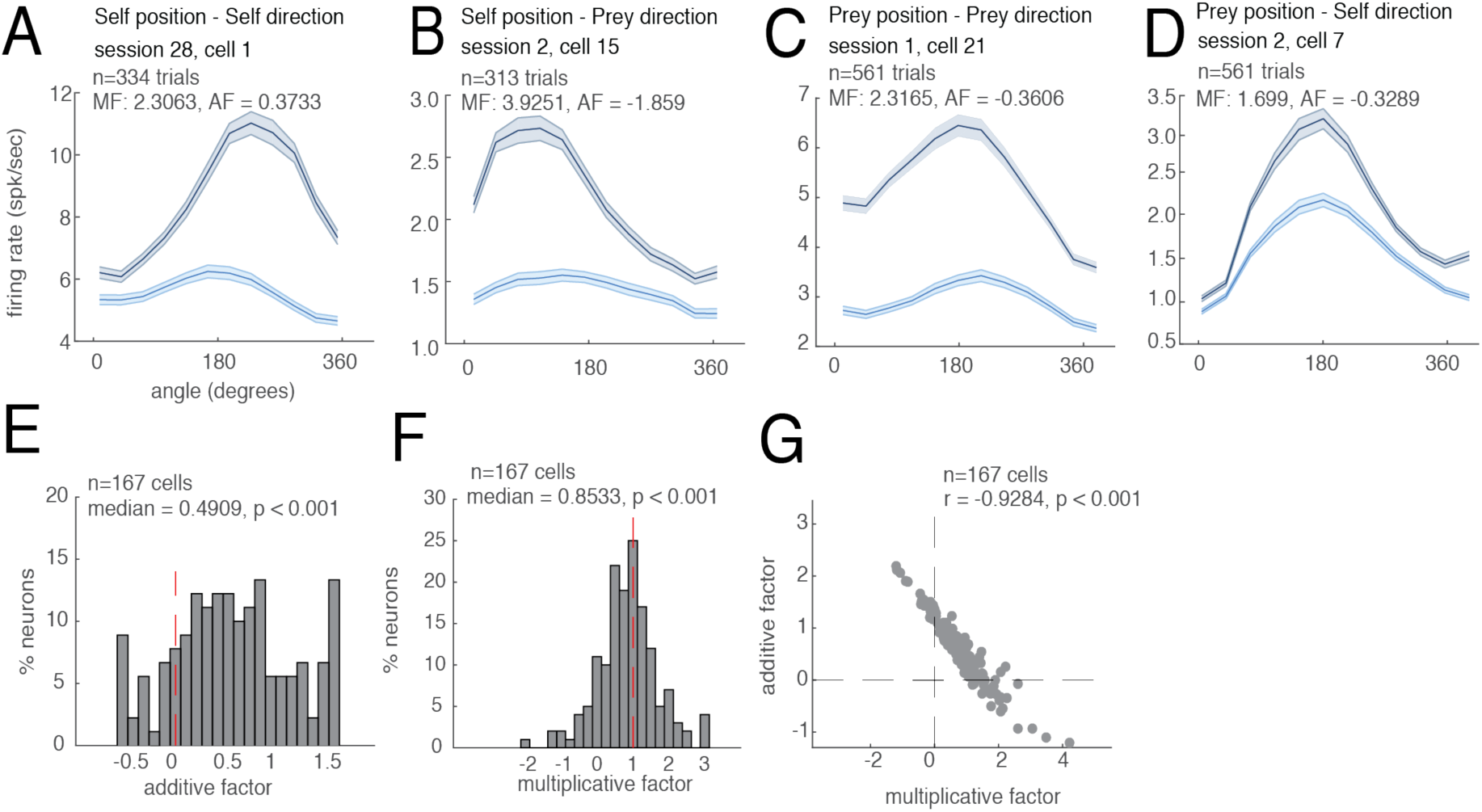
Mixed Selectivity and Adaptivity. We illustrate our findings on mixed selectivity by focusing on relative angle. See supplement for other variables. **(A-D)** Angular tuning curves for example neurons that showed significant mixed selectivity. Horizontal axes reflect angle between self and prey; 0 indicates heading towards the prey. Neurons show angular selectivity that is modulated by anticipated reward. Gray line: high-value prey. Blue line: low-value prey. (A) multiplicative influence of self-direction by self-position, (B) self-direction by prey-position, (C) prey-direction by self-position, (D) prey-direction by prey-position. **(E-F)** Additive (E) and Multiplicative (F) Factors of population. AF>zero indicates additive interaction; MF deviates from 1 indicates multiplicative interaction. **(G)** Relationship between MF and AF. Negative correlation indicates that neurons with multiplicative interaction are less likely to have additive interaction.

We confirmed this mixed selectivity result with an additional method that is less sensitive to the shape of the tuning curve [32]. Specifically, we characterized the range (max firing rate – min firing rate) of each tuning curve as a function of the mean firing rate for the position (three bins; method from ref 32). As expected under mixed-selectivity, the range increased with mean position segment firing rate (median r=0.2305, p<0.001, rank-sum test; Figure S3). Together these two results indicate that dACC neurons use nonlinearly mixed selectivity (and not just multiplexing) to encode various movement-related variables.

### Spatial coding is distributed across neurons

Although responses of a large number of neurons are selective for spatial information about the three agents, it is not clear to what extent a broad population drives behavior [49]. Thus, we examined how much each neuron in the population contributes to behavior using population decoding with an additive method [50]. If only small sets of neurons contribute to behavior, the decoding performance with respect to number of neurons will soon reach a plateau. We randomly assigned neurons to the decoder regardless of whether they were significantly tuned to the variable of interest. We found that as the number of neurons included for decoding analysis increases, the accuracy of decoding positional variable (both self and prey) increases without evidence of saturation (Figure 7A).

**Figure 7.**
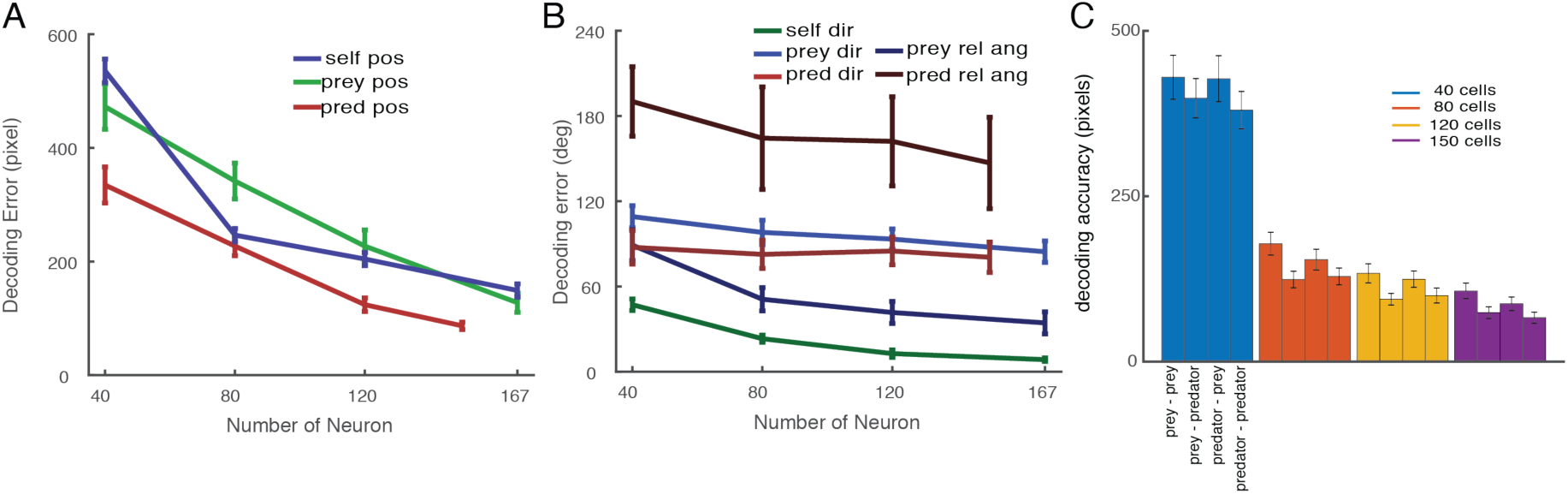
**(A)** Population decoding accuracy (distance between model and empirical data) for self (blue) position, prey (green), and predator (red) position improves (i.e. gets lower) as number of neurons analyzed rises. This analysis indicates that these variables are encoded in a distributed manner, and that all three agents can be readily decoded. **(B)** Population decoding accuracy for self (dark green) direction, prey (purple) direction, predator (red) direction, prey relative angle (dark blue) and predator relative angle (maroon). All curves decrease modestly, indicative of distributed decoding. (**C)** Time-to-time decoding performance when either prey is closer or predator is closer. Then the decoding performance for either prey position or predator position was estimated. Decoding accuracy was estimated as a function of number of cells, in groups corresponding to 40, 80, 120, and 150 cells.

We next applied this serial decoding procedure to examine relative strength of different formats of spatial coding. For this analysis, we focused on coding of world-centric angle (self- and prey-direction) and avatar-centric angle, which share common units (specifically, degrees; Figure 7B). We find that the strength of information within the neural population is mixed between world-centric and avatar-centric information. Self-direction information is strongest and prey direction information is weakest (and their difference is significant, decoding error by using all neurons are 8.496±0.603°, 34.364±3.795°, 84.455±3.712°, p< 0.001, ANOVA). This result showing distributed information contrasts with previous findings in a similar paradigm that show positional variables are encoded by only a handful of neurons [49]. We speculate that difference may due to the complexity of our task, which may require a high-dimensional neural space to maximize the information [51,52].

### Reward encoding

Research based on conventional choice tasks indicates that dACC neurons track values of potential rewards [53]. We next asked how dACC encodes anticipated rewards in our more complex task. Initially, we regressed reward variable against the neural activity one second before the trial end for all types of trial. We found that, averaging over all other variables, the value of the pursued reward modulates activity of 9.3% of neurons (n=14/150, p=0.0227, one-way binomial test). Note that this analysis ignores the potential encoding of prey speed, which is perfectly correlated with static reward in our task design. We then explored possibility of reward being modulatory variable, which means that reward increase the other variables’ selectivity. We find that tuning for all variables increases with increasing reward (p<0.05 in each case, sign-rank test, Figure 8). Compare to the random split of data, which yielded insignificance difference in tuning, splitting data according to the value of prey did yield a significant difference in tuning proportions for the variables (Figure 8). Importantly, the percent of neurons tuned for each variable is maintained in the random split, indicate reliability of tuning. Instead, the proportion of neurons whose responses were selective for self-position was not different when the data was split randomly into half (28.0 % vs. 27.4%, p=0.2783, sign rank test, 50 times bootstrapping).

**Figure 8.**
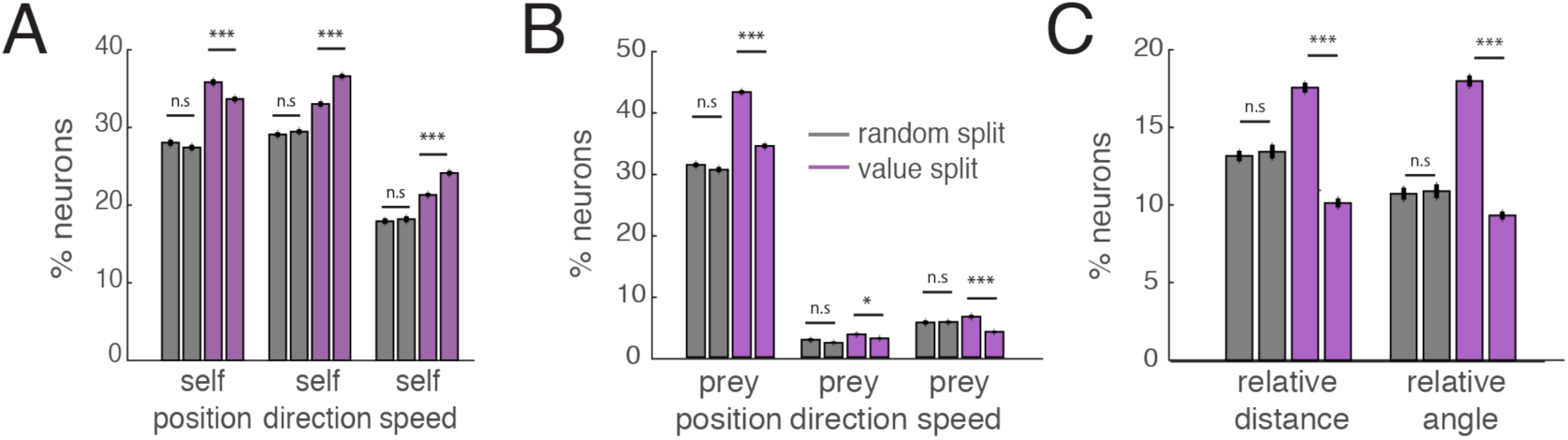
Tuning for task variables change with prospective reward. These data support the idea of mixed selectivity. **(A)** The number of tuned neurons for self-variables (self-position, self-direction, and self-speed) when splitting data randomly (grey bar) or according to value of pursued prey (purple bar). Splitting by value increases proportion of tuned neurons, indicating that value modulates responding in a systematic way. **(B)** The number of significantly tuned neurons for prey variables (prey position, prey direction, and prey speed) when splitting data randomly (grey bar) or according to value of pursued prey (purple bar). The difference of value split was significant (p = 0.0221 for prey speed, and p < 0.001 for other prey variables) but not for random split. **(C)** The number of significantly tuned neurons for egocentric (self-position, self-direction, and self-speed) when splitting data randomly (grey bar) or according to value of pursued prey (purple bar). The difference of value split was significant (p < 0.001).

### Gaze does not change selectivity of spatial tuning

Activity in dACC is selective for saccadic direction and may therefore also correlate with gaze direction [54]. Consequently, it is possible that our spatial kernels may reflect not task state but gaze information. In fact, the apparent selectivity of dACC neurons for each agent’s position could artifactually result from gaze tuning if the subject periodically fixated each target. To resolve the confound, we repeated our GLM analyses but included eye position (only for the one subject from which we collected gaze data). We found that that the number of tuned neurons for the position of any agent did not substantially change; that is, that adding in gaze position as a regressor did not qualitatively change our results (Figure 9).

**Figure 9.**
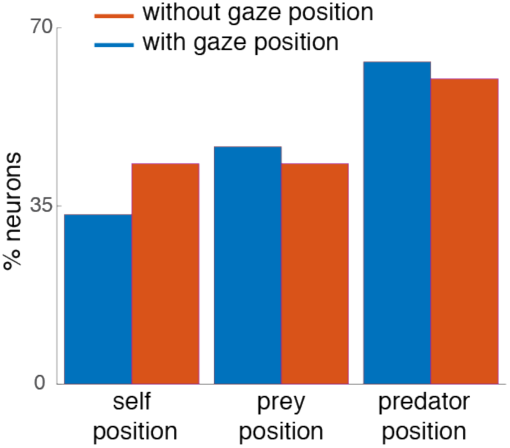
Analysis to detect potential gaze confounds. Red bars: proportion of neurons tuned for three key world-centric variables using the standard GLM described above. Blue bars: same results, but this time from a version of the GLM that included eye position as a regressor. That version is constructed so that all variance possible associated with eye position is assigned to eye position first and only residual encoding of task variables is counted towards those variables. All three variables are still significantly observed in the population when including gaze position.

## DISCUSSION

We examined the neural foundations of pursuit and evasion by recording single unit activity in the dACC while rhesus macaques performed a joystick-controlled pursuit task (Figure 1). We find that dACC carries a dynamic (i.e. continuously updated) multi-centric (i.e. both world-centric and avatar-centric) multi-agent (self, prey, and predator) representation of the state of the task. These results indicate a clear role of the dACC in mapping functions that are intrinsic to pursuit. One limitation of the present study is that we did not record activity in other regions that may also be involved. Therefore, we cannot conclude that dACC plays a unique role in this process. Future studies will be needed to functionally differentiate dACC from other regions.

What is the benefit of encoding both absolute (world-centric) and relative (avatar-centric) maps? One possibility is that dACC participates in the process of mediating between the two representations. Another (not mutually exclusive) possibility is that both representations are important for behavior. Consider, for example, that avatar-centric codes may allow for rapid on-the-fly changes in trajectory while world-centric ones may allow for more abstract planning, for example, allowing the subject to trap the prey in corners. Having both in the same place may allow for their coordination to make optimal decisions. Indeed, this idea is consistent with the idea that a major function of dACC is to use multiple sources of information to set and drive a strategy from a high vantage point [23–25,41,43].

Neuroscientists are just beginning to understand the neural basis of tracking of other agents. Traditionally, neurons in primate dACC and its putative rodent homologues are not expected to encode place fields. For example, a putative rodent homologue is reported to utilize positional information but not signal place per se [27]. However, at least one notable recent study has demonstrated that place field information can be decoded from rodent ACC [49]. Our results build on this finding and extend our understanding of the spatial selectivity of this region further to tracking of other agents. Two recent studies demonstrate the existence of coding for positions of conspecifics in the CA1 region of the hippocampus in rats and bats [55,56]. Our results here extend on them in three ways. First, they confirm speculation that positional tracking extends to at least one hippocampal target region in the prefrontal cortex. Second, they demonstrate that positional tracking extends to multiple agents, including different types (prey and predators), and that it is multicentric. Most intriguingly, and most speculatively, our results directly link tracking of others to personal goal selection processes.

Although the responses we observe have some similarity to hippocampal place cell firing, dACC responses are less narrowly localized than place cells, are less patterned than grid cells, and can only be detected using a newly developed statistical approach [32]. While the medial entorhinal cortex is commonly associated with grid cells [57–59], one recent study demonstrated that it carries a much richer set of spatial representations [32]. Our study indicates that such non-canonical spatially-mapped neurons are not limited to entorhinal cortex, or to rodents, and can be observed in virtual/computerized environments, and extend to other agents in the environment. These results confirm the highly embodied role of dACC in economic choice and highlight the central role of spatial information in economic decision-making [10,11,60–62].

Our data are limited to a single region and do not imply a unique role for this region. Most notably, several other ostensibly neuroeconomic brain regions carry rich spatial repertoires, including OFC [63] and vmPFC [64,65]. These regions also have connectivity that includes, directly or indirectly, medial temporal navigation regions and motor and premotor regions. Therefore, we predict that the patterns we observe here would also be observed, albeit perhaps more weakly, in these other regions. Unlike these regions, the dACC has been linked to motor functions, albeit much less directly than, for example, motor cortex [24]. What is new here, then, is the observation that dACC tracks the kinematics of self, prey, and predator, uses multicentric tuning for these multiple agents. In addition to what it tells us about dACC, the multi-centric multi-agent tuning also serves as a control for possible motor effects explaining the results.

## Conclusion

Most studies of the neural basis of decision-making focus on simple and abstract choices but natural decisions take place in a richer and more complex world. In our task, decisions are continuous – they take place in an extended time domain and the effects of decisions are manifest immediately. Moreover, our task, and monkeys’ ability to perform it well, illustrate the complexity of the word decision – it has a simple and clear definition in economic choice tasks. But in a more naturalistic context, like this one, it can refer either to the specific direction the subject is moving at a point in time, or to the higher level goal of the subject. Ultimately, we anticipate that consideration of more complex tasks may lead to a refinement of the concept of decision.

More broadly, given the critical role of foraging in shaping our behavioral repertoires overall [66–68], we and others have proposed that spatial representations are likely to be a ubiquitous feature of our reward and decision-making systems [61,69]. This idea is supported, at the most basic level, by studies showing clear spatial selectivity in the reward system in both rodents and primates [63,64,70–73]. In other words, spatial information is not abstracted away even in ostensibly value-specialized regions [61,74]. By utilizing ever more complex paradigms, we can place the brain into natural states not probed by conventional tasks and uncover unanticipated complexities in neuronal responses.

## Acknowledgements

We thank Alex Thomé for his critical role in designing the task, for devising the training protocols, and for developing our joysticks. We thank Marc Mancarella for his critical help with joystick training. We are grateful for helpful discussions from Habiba Azab, Steve Piantadosi, Marc Schieber, and Adam Rouse.

## STAR Method

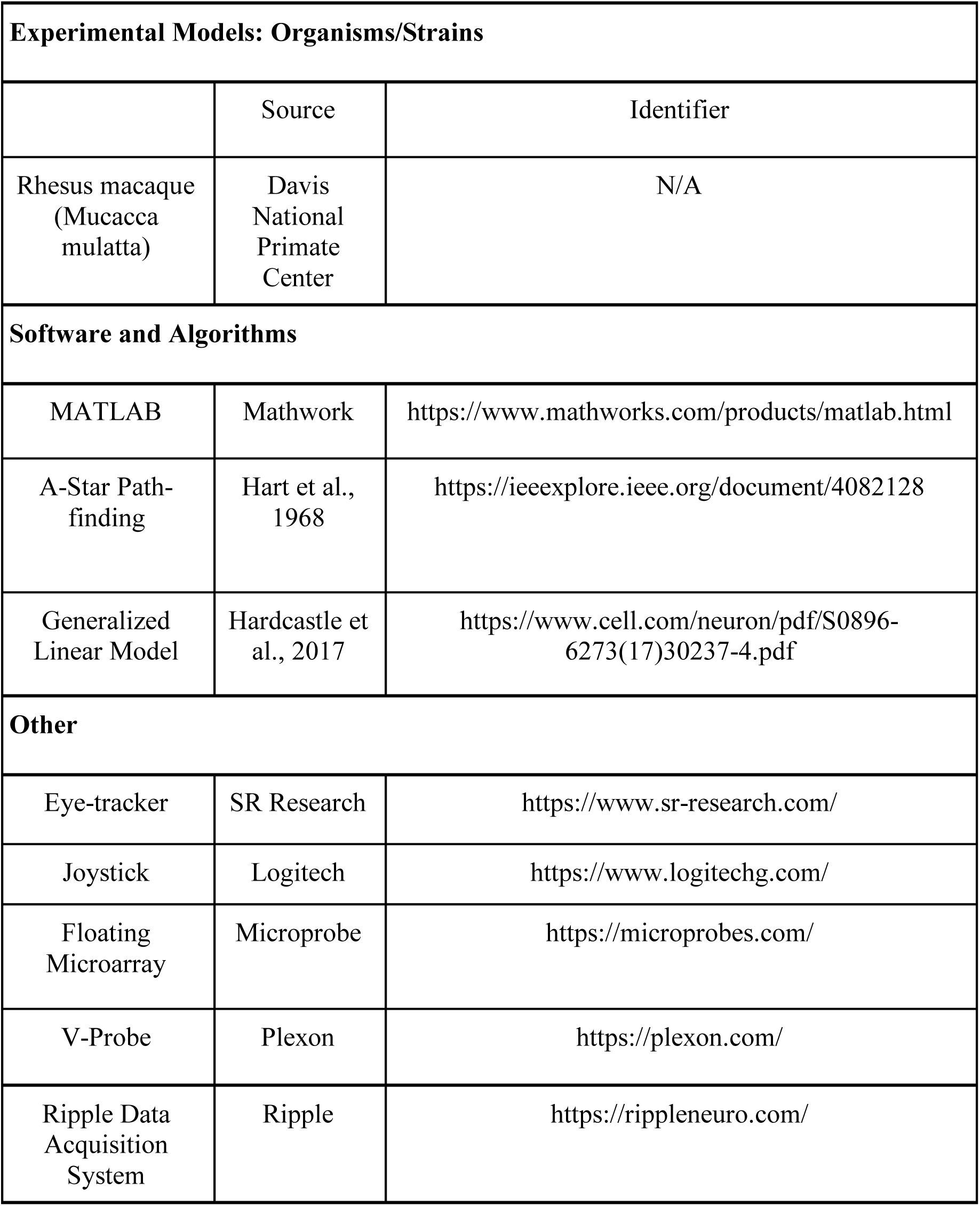

## EXPERIMENTAL METHODS

All animal procedures were approved by the University Committee on Animal Resources at the University of Rochester or the Institutional Animal Care and Use Committee (IACUC) at the University of Minnesota. All experiments were designed and conducted in compliance with the Public Health Service’s Guide for the Care and Use of Animals.

### Subjects

Two male rhesus macaques (*Macaca mulatta*) served as subjects. Note that the inclusion of only two subjects (as is standard in primate physiology) does reduce our ability to draw inferences about group effects. Previous training history for these subjects included a variant of a Wisconsin card-sorting task (subject H, 74), basic decision-making tasks (subject H, 75), and two foraging tasks (both subjects, 76–78).

### Experimental Apparatus

The joystick was a modified version of a commercially available joystick with a built-in potentiometer (Logitech Extreme Pro 3D). The control bar was removed and replaced with a control stick (15 cm, plastic dowel) topped with a 2’’ plastic sphere, which was custom designed through trial and error to be ergonomically easy for macaques to manipulate. The joystick position was read out in MATLAB running on the stimulus-control computer.

### Task Design

At the beginning of each trial, two or three shapes appeared on a gray computer monitor placed directly in front of the macaque subject (1920×1080 resolution). The yellow (subject K) or purple (subject H) circle (15-pixel diameter) was an avatar for the subject and moved with joystick position. A square shape (30-pixel length) represented the prey. The movement of the prey determined by a simple algorithm (see below). Successful capture is defined as any overlap between the avatar circle and the prey square. Each trial ends with either successful capture of the prey or after 20 seconds, whichever comes first. Capture results in immediate juice reward; juice amount corresponds to prey color: orange (0.3 mL), blue (0.4 mL), green (0.5 mL), violet (0.6 mL), and cyan (0.7 mL). Failure to capture results in timeout and a new trial. (Failures were rare).

The path of the prey was computed interactively using A-star pathfinding methods, which are commonly used in video gaming [80]. For every frame (16.67 ms), we computed the cost of 15 possible future positions the prey could move to in the next time-step. These 15 positions were spaced equally on the circumference of a circle centered on the prey’s current position, with radius equal to the maximum distance the prey could travel within one time-step. The cost in turn is computed based on two factors: the position in the field and the position of the subject’s avatar. The field that the prey moves in has a built-in bias for cost, which makes the prey more likely to move towards the center (Figure 1A). The cost due to distance from the subject’s avatar is transformed using a sigmoidal function: the cost becomes zero beyond a certain distance so that the prey does not move, and the cost becomes greater as distance from the subject’s avatar decreases. the position with the lowest cost is selected for the next movement. If the next movement is beyond the screen range, then the position with the second lowest cost is selected, and so on.

The maximum speed of the subject was set to be 23 pixels per frame (i.e. 16.67 ms). The maximum and minimum speeds of the prey varied across subjects and were set by the experimenter to obtain a large number of trials (Figure 1). Specifically, speeds were selected so that subjects could capture prey on above 85% of trials; these values were modified using a staircase method. If subjects missed the prey three times consecutively, then the speed of the all prey was reduced temporarily. The minimum initial distance between the subject avatar and prey was 400 pixels. The strict correlation between speed and value means that value cannot be directly deconfounded in this study.

A predator (triangle shape) appeared on 25% of trials. Capture by the predator led to a time-out. Predators came in five different types (indicated by color) indicating different level of punishment, ranging from 2 seconds to 10 seconds. The algorithm of the predator is to minimize the distance between itself and player. Unlike the prey, the predator algorithm is governed by this single rule.

The design of the task reflects primarily the desire to have a rich and variegated virtual world with opportunities for choices at multiple levels that is neither trivially simple nor overly complex. The decision to include a condition with multiple prey was added specifically for these reasons and for the additional reason that we wanted to verify that subjects distinguished the differently valued prey by pursuing them with differential preference.

The reason we deliberately confounded reward and speed was to make sure the task neither too difficult nor too easy, and to ensure that the results of the animals’ choices between prey were interesting and meaningful. We also wanted to keep the effort/interest level roughly the same on each trial.

### Trajectory-Based Trial Sorting

In 50% of trials, subjects saw two prey items instead of one. We developed the TBTS (Trajectory-Based Trial Sorting) method to determine which prey the subject was pursuing at any given time. This method requires to calculate 1) the angle differences between subject’s and each prey’s trajectory from time *t-1* to *t*, 2) change of distance between subject and each prey (estimate whether prey is getting closer to subject or floating) and 3) dynamic time warping outcome (to calculate the distance between the signal) between the trajectory of the subject and each prey. Then we multiply them to obtain a single scalar for each agent at every time point, and then smoothed with a boxcar of 5 frames to make secure autocorrelation between the data points. The prey being pursued will have smaller angle difference with the subject, the distance between the subject and pursued prey will be decreasing (due to avoiding algorithm, non-chased prey will tend to increase its distance with the subject), and dynamic time wrapping outcome will be smaller. Thus, when one prey is pursued continuously, then this value will stay always smaller than the other. From there, we excluded trials with switches to avoid any confounds that arose from not knowing what prey the subjects are pursuing (about 5% of trials overall).

### Electrophysiological recording

One subject was implanted with multiple floating microelectrode arrays (FMAs, Microprobes for Life Sciences, Gaithersburg, MD) in the dorsal anterior cingulate cortex (dACC). Each FMA had 32 electrodes (impedance 0.5 MOhm, 70% Pt, 30% Ir) of various lengths to reach different dACC. Neurons from another subject were recorded with laminar V-probe (Plexon, Inc, Dallas, TX) that had 24 contact points with 150 µm inter-contact distance. Continuous, wideband neural signals were amplified, digitized at 40 kHz and stored using the Grapevine Data Acquisition System (Ripple, Inc., Salt Lake City, UT). Spike sorting was done manually offline (Plexon Offline Sorter). Spike sorting was performed blind to any experimental conditions to avoid bias.

### Tracking Neurons Over Multiple Days

We used an open-source MATLAB package “Tracking Neurons Over Multiple Days” [81]. Briefly, pairwise cross-correlograms, the autocorrelogram, waveform shape, and mean firing rate were used together as identifying features of a neuron. For classifying the identical neurons across the session, we calculated four values that characterize individual neuron: mean firing rate, autocorrelation, pairwise correlation with other neurons, and shape of the waveform. Then we applied a quadratic classifier that computes an optimal decision boundary under the assumption that the underlying data can be modeled as a mixture of multivariate Gaussians [81].

### Details of LN model

To test the selectivity of neurons for various experimental variables, we adapted Linear-nonlinear Poisson Models (LN models). The LN models estimated the spike rate (r*_i_*) of one neuron during time bin t as an exponential function of the sum of the relevant value of each variable at time t projected onto a corresponding set of parameters (w*_i_*). The LN models can be expressed as:

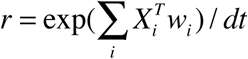

Where r denotes a vector of firing rates for one neuron over T time points, *i* indexes the variables of interest, e.g. position of avatar on screen. X*_i_* is a matrix where each column represents an “state” of the animal (e.g. one of twelve speeds, determined by post-hoc binning) obtained from binning the continuous variable so that all the columns for a particular row is 0 except for one column (one-hot encoding).

Unlike conventional tuning curve analysis, GLM analysis does not assume the parametric shape of the tuning curve *a priori*. Instead, the parameter weights, which defines the shape of tuning for each neuron, were optimized by maximizing the Poisson log-likelihood of the observed spike train given the model expected spike number (n), with additional regularization for the smoothness for parameters in a continuous variable and a lasso regularization for parameters in a discrete variable. Position parameters are smoothed across rows and columns separately. The regularization hyperparameter was chosen with maximizing cross-validation log-likelihood based on several randomly selected neurons. The optimization was performed with a MATLAB built-in function (fminunc). Model performance of each neuron is quantified by the log-likelihood of held out data under the model. This cross-validation procedure was repeated 10 times and overfitting was penalized. Thus, we can compare performance of models with varying complexity.

### Forward model selection

Model selection was based on the cross-validated log-likelihood value for each model. We first fitted n models with a single variable, where n is the total number of variables. The best single model was determined by the largest increase in spike-normalized log-likelihood from the null model (i.e., the model with a single parameter representing the mean firing rate: r). Then, additional variables (n-1 in total) were added to the best single model. The best double model was preferred over the single model only if it significantly improves the cross-validation log-likelihood (Wilcoxon Signed Rank Test, α = 0.05). Likewise, the procedure was continued for the three-variable model and beyond if adding more variables significantly improved model performance, and the best simplest model was selected. The cell was determined to be not tuned to any of the variables considered if the log-likelihood increase was not significantly higher than baseline.

### Response profile

We derive response profiles from filter of model for a given variable j to be analogous to a tuning curve of given variable. These were computed as, which α =Πi [_i is all other variables than j_]mean(exp(w_i_)) is a scaling factor that marginalizes out the effect of the other variables. The *dt* transforms the units from bins to seconds. Thus, for each experimental variable, the exponential of the parameter vector that converts animal state vectors into firing rate contributions is proportional to a response profile; it is a function across all bins for that variable and is analogous to a tuning curve.

### Principal Component Analysis (PCA) and clustering relative distance tuning

We reasoned that relative distance between subject and prey is encoded and is not simply an artifact of proximity to reward acquisition. This variable is encoded in dACC, although generally with robustly positive and monotonic code [42,77,82]. Instead, complex shape of tuning may indicate distance is encoded. To examine this, we clustered the tuning curves according to shape to show whether there might be some functional clusters. First, we selected out 60 neurons that are individually significantly tuned to relative distance of prey. Then, we performed dimensionality reduction via PCA and found 2 PCs explain 70% of variance in the data. Thus, we projected data into first two PCs and performed k-nearest neighbor and found elbow with K=4 (Figure 5D). Identical method was used for profiling the filters that are tuned for speed (Figure 4).

### Multiplicative vs. Additive shift of tuning

We report that neurons exhibit ‘multiplicative’ tuning, defined as r(x, y) =r(x)* r(y), which means the tunings for each variable interact non-linearly, and thus have mixed selectivity [48]. However, there is possibility that the neurons might show additive tuning, defined as r(x, y) =r(x)+ r(y). Strictly, linear addition would be multiplexed but not mixed selectivity [45].

Differentiating between these two has important implications as multiplicative coding may point to a fundamental transformation of information, while additive coding suggests signals simply linearly combine [32,48].To quantify the nature of conjunctive coding and verify our assumption that tuning curves multiply, we examined neurons that significantly encoded both position of two agents (self and prey) and direction of two agents based on model performance (e.g., both the position and other models had to perform significantly better than a mean firing rate model). We examined differences in how the tuning curve for specific variables r(x, y), or the tuning curve across y for a fixed value x*, will change as a function of r(x*) to estimate whether neurons exhibit multiplicative or additive. In the multiplicative model, an variation of r(x*) will modify the shape of tuning curve (either stretch or compress) r(x*, y), whereas in the additive model it will shift r(x*, y) simply up or down. To quantify these differences, we took x to be position and y to be either direction or speed of agent, and binned position into 15×15 bins. We then calculated the firing rate for each position bin (i.e., computed r(x*) for every x*), sorted the position bins according to firing rate values, and divided the bins into two (high vs. low, for analysis 1. see below) or three (for analysis 2, Figure S3) segments. Each segment, thus, corresponded to a location of the environment with approximately the same firing rate. We then generated a series of tuning curves (either direction or speed) based on the spikes and directions visited during each segment.

Once we obtained tuning curves for each segment, for each single neuron, we characterized its multiplicative, additive, or displacement modulation with population activity by performing linear regression on the average response to each state bins, when population activity was high compared with when it was low. The slope of the linear fit indicates how tuning scales multiplicatively with population activity (so called, multiplicative factor [MF]). The slope deviating from 1 shows either multiplicative or displacement interaction. The intercept of the fit describes the additive shift to tuning with population activity. To obtain a relative measure of the additive shift, like the MF, we defined the additive factor (AF) as the ratio between this intercept and the mean firing rate of the neuron averaged.

We additionally confirmed the multiplicative tuning shift by computing the range (maximum firing rate – minimum firing rate) of tuning curve as a function of the mean firing rate for position segment *i*. If each neuron shows multiplicative tuning shift, the range should increase with position segment. Whereas the additive model result in constant range. The range of the tuning curve and mean position segment firing rate exhibited a positive slope in pool of significantly tuned neurons for self direction, self speed, prey direction and prey speed (134/165 pooled neurons; median slope > 0 with p < 0.001, two-way binomial test).

### Downsampling for Reward Modulation Analysis

Each session was split into low vs. high reward of pursued target. To match the coverage of experimental variable for each condition, we first binned position, direction, speed into 225 bins, 12 bins, and 12 bins, respectively, and computed the occupancy time for each bin. The coverage of experimental variables across conditions was matched by down-sampling data points from either condition so that occupancy time was matched for each bin, with points removed based on the difference in direction occupancy. For this analysis we wanted to have the total number of spikes match in the compared conditions. We did this because spikes convey information and we did not want to introduce spurious differences in information content that reflected only random variation in number of spikes. We then repeated this procedure 50 times and took the average result. We did this because the decimation procedure is stochastic and there susceptible to random effects. We can reduce these effects and obtain a more precise estimate of the true effect, through repetition and averaging.

### Decoding Analysis

Decoding accuracy was assessed by simulating spike trains, which were based on Poisson statistics and a time-varying rate parameter. In each group, spikes (n_c_) for neuron c were generated by drawing from a Poisson process with rate r_c_; where (w_i,c_ are the learned filter parameters from the selected model for neuron c, X_i_ is the behavioral state vector, and i denotes the experimental variables). If the model selection procedure determined that a neuron did not significantly encode variable i, then w_i,c_ = 0. Next, the simulated spikes were used to estimate the most likely variable that is being decoded. To decode experimental variables at each time point t under each decoder, we estimated the animal state that maximized the summed log-likelihood of the observed simulated spikes from t-L to t:

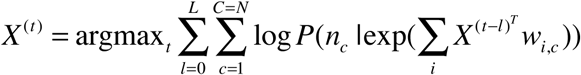

where C is the number of cells in that population. Decoding was performed on 50 randomly selected 2000 ms (L= 121 time bins) of session. The average position decoding error (pixel distance error), direction (error in degree), and speed (error in pixel per second) were recorded. For examining the coding scheme of the population (whether sparse or distributed code), we increased number of neurons being included in this analysis from 40 to 167 by 40 neurons. The random shuffling of neurons was performed 30 times.

### Adaptive Smoothing Method

An adaptive smoothing method is used for presentation purposes although not for quantified data analysis [83]. Briefly, the data were first binned into 100×1 vector of angle bins covering the whole 360 degrees of the field, and then the firing rate at each point in this vector was calculated by expanding a circle around the point until the following criteria was met:

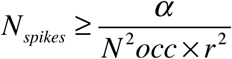

where N_occ_ is the number of occupancy samples, N_spikes_ is the number of spikes emitted within the circle, r is the radius of the circle in bins, and alpha is a scaling parameter set to be 10000 as previous studies.

## Supplementary Material

### Figure Caption for Supplementary Figures

**Figure. S1.**
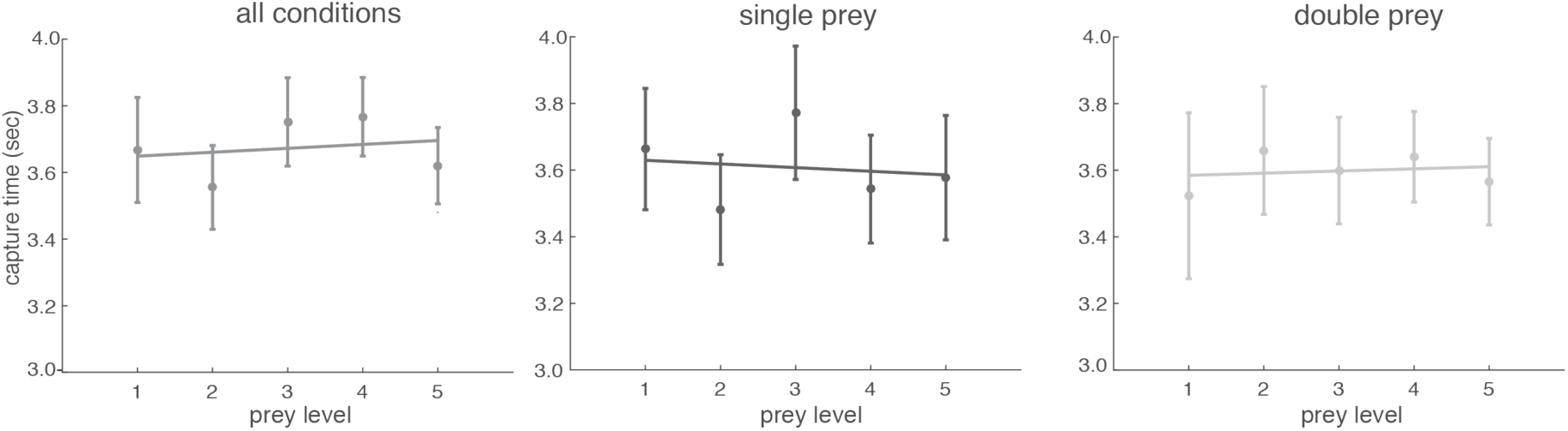
Flattened capture time regardless of task conditions. We investigated individual conditions for capture time and found that the capture time does not differ among the prey difficulty when it is single prey or double prey.

**Figure. S2.**
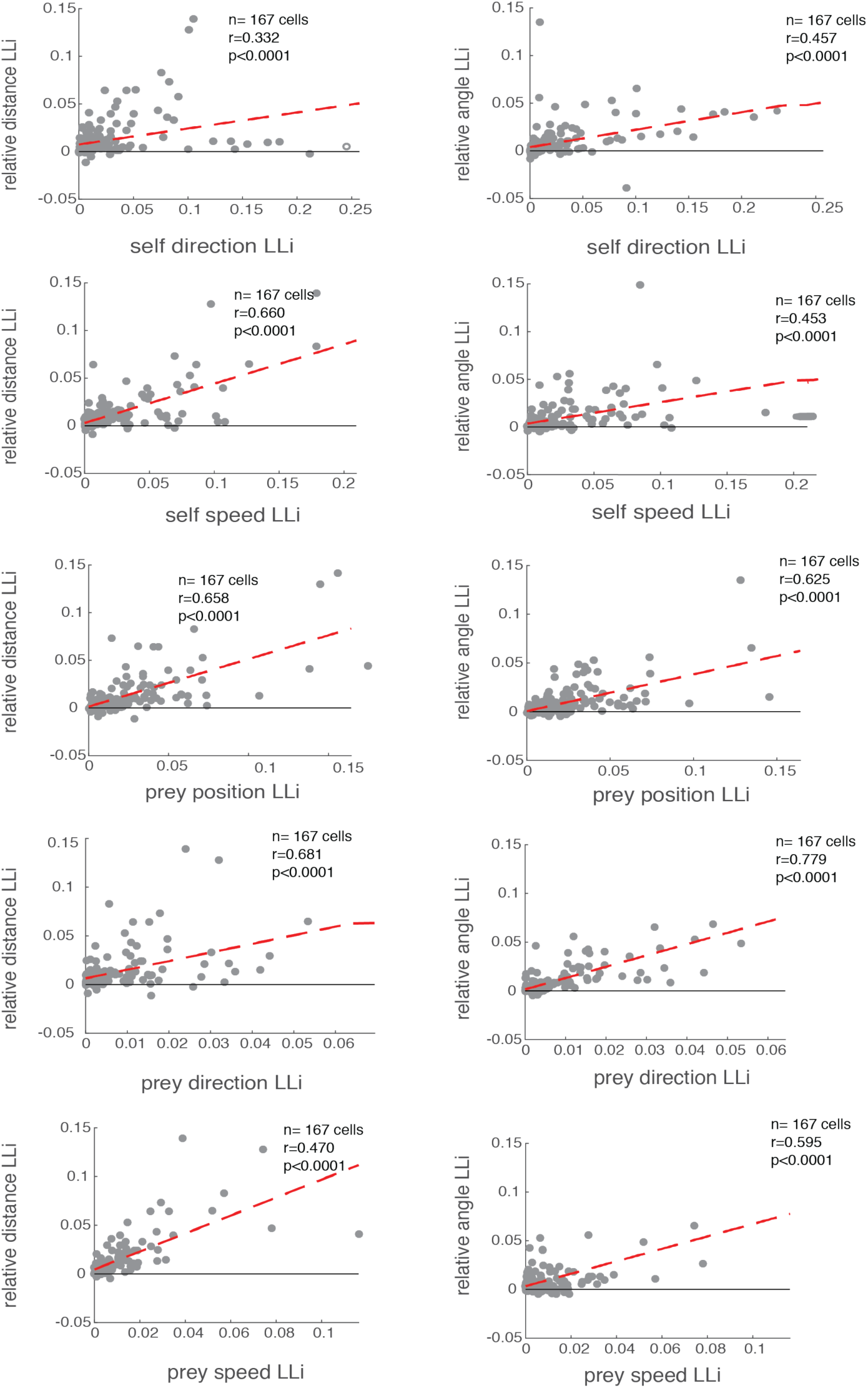
Log-Likelihood comparison between multiple variables. Scatter plots illustrating correlation between log-likelihood increase (LLi) for various experimental variables. Each dot corresponds to one neuron. Red line indicates the line obtained from linear regression (r-value and p-value is at each panel).

**Figure. S3.**
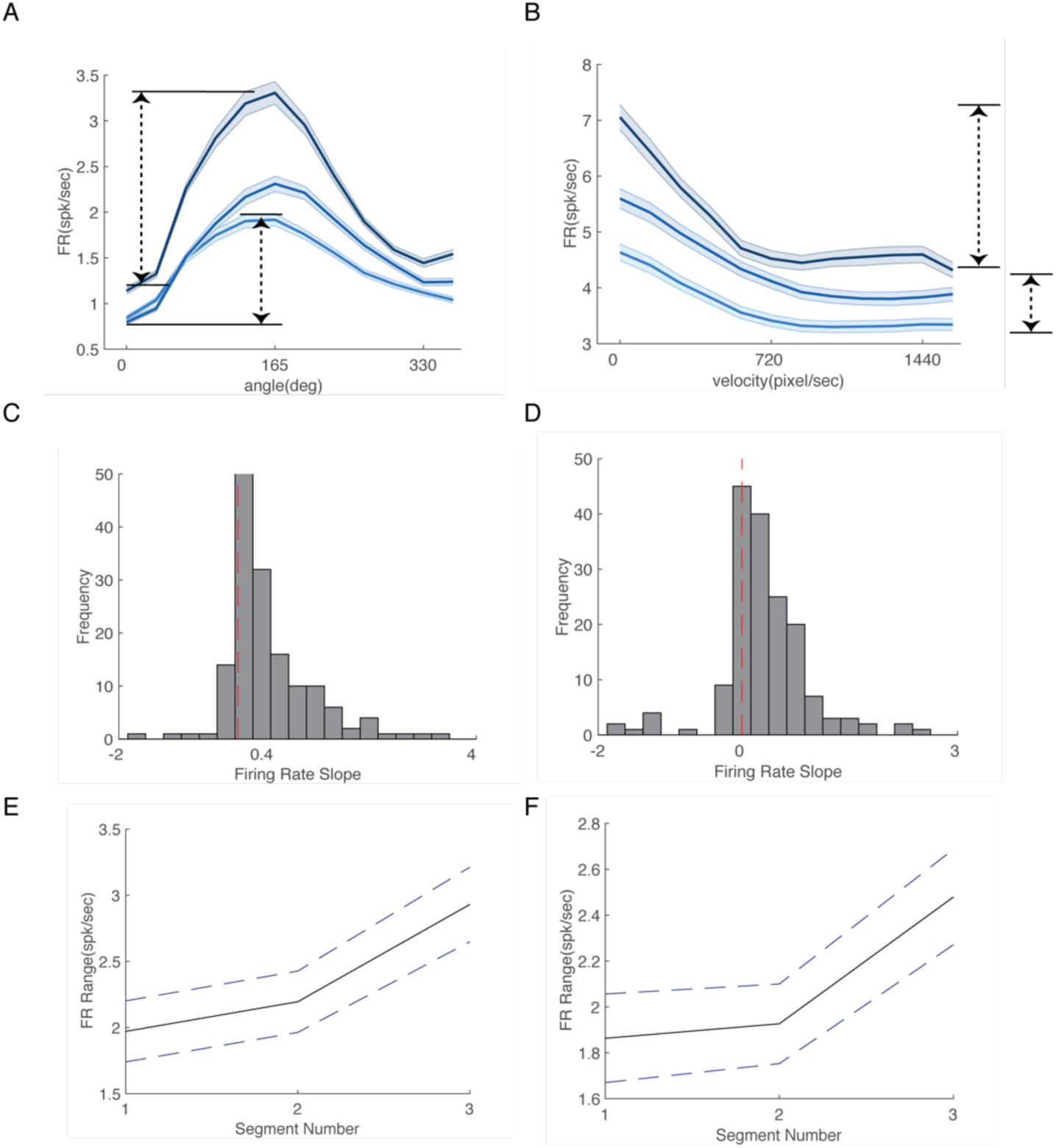
Additional method for examining mixed-selectivity. **(A-B)** The individual neurons that has unimodal tuning curve (left) and non-unimodal tuning curve (right). Tuning curve shape variability could make first method used for mixed-selectivity detection less sensitive (the right one has barely significant MF, 1.0194). Thus, adopting method used in Hardcastle et al., 2017, we measured the firing rate angle in each condition and compared them. If the slope is positive when each condition is plotted against range of firing rate, then it is considered as mixed-selective. **(C)** The slope of pooled result for position influence to direction, which is significantly different from zero (slope = 0.4798, p < 0.001). **(D)** The slope of pooled result for position influence on speed, which is significantly different from zero (slope = 0.3080, p < 0.001). **(E-F)** The mean and standard error of the mean for the firing range at each condition. It shows increasing trend.

